# Global Lipidomic Analysis of Lytic KSHV Infection: The lipid chaperone FABP4 is required for maximal infectious virion production

**DOI:** 10.64898/2026.04.19.719449

**Authors:** Eranda Berisha, Erica L. Sanchez

## Abstract

Kaposi’s Sarcoma Herpesvirus (KSHV), an enveloped double-stranded DNA virus, is the etiological agent of Kaposi’s sarcoma (KS), an endothelial cell-based tumor. KSHV is a leading cause of infection-related cancers in sub-Saharan Africa and immunocompromised individuals worldwide. Therefore, it is vital to identify the underlying mechanisms of viral infection and transmission to effectively identify specific therapeutic strategies and combat the disease. Here, we demonstrate that KSHV rewires the host cell lipidome during lytic infection. Bulk lipidomic analysis shows significant changes in the abundance of neutral lipids and phospholipids during lytic infection. We further investigated fatty acid-binding proteins (FABPs) to understand the underlying mechanisms that support KSHV pathogenesis. Using the doxycyclin-inducible iSLK.BAC16 cell line, we find that FABP genes are differentially regulated by lytic KSHV infection compared to latent infection. We report that FABP4 is significantly upregulated during lytic infection. Loss of FABP4 during lytic infection does not impact viral gene transcription however, lytic protein translation is reduced. Moreover, our intracellular and extracellular viral titers indicate that FABP4 affects maximal infectious virion production. This study highlights the role of FABP4 and its therapeutic potential as a target that facilitates KSHV infection and pathogenesis.

## Introduction

Kaposi’s Sarcoma (KS) is an endothelial cell-based tumor endemic to sub-Saharan and the Mediterranean, and the most common cancer in HIV-infected individuals[1–3]. Kaposi’s Sarcoma-Associated Herpesvirus (KSHV), the causative agent of KS, is an oncogenic enveloped double-stranded DNA virus and member of the *Gammaherpevirinae* subfamily. KSHV infection induces several gene expression and metabolic changes in the host that facilitate maximal virion production. KSHV has both latent and lytic viral phases, with ∼95% of KS tumor cells undergoing latent infection, and only 1-5% cells lytically replicating[1–4]. The latent phase of KSHV is marked by limited viral gene expression, no virion production, and host cell survival. However, upon reactivation, all viral genes are expressed, and virion production occurs, eventually leading to host cell lysis[4,5]. Ultimately, both the latent and lytic phases are essential for KSHV pathogenesis.

Previous publications have reported significant alterations in host cell metabolism, specifically lipid metabolism, that occur during the latent phase of KSHV infection in endothelial cells[6–9]. Latent KSHV infection induces *de novo* lipogenesis and requires fatty acid synthesis for survival. Furthermore, treatment with fatty acid synthesis inhibitors induces apoptosis of latently infected cells[7]. An interplay between increased lipogenesis and lipid droplet formation, cytoplasmic organelles that store neutral lipids, occurs during latent infection[6–8]. Another study has reported that latent infection induces an increase of cholesterol ester (CE) synthesis and CE content in lipid droplets for membrane biogenesis[9]. However, lipid droplet degradation through either lipolysis or lipophagy generates free fatty acids that can be used to produce ATP through mitochondrial ꞵ-oxidation or be utilized for peroxisomal ꞵ-oxidation[6,8,9]. Interestingly, there are limited reports of lipidomic alterations during KSHV lytic infection.

The lytic cascade and the distinct stages of lytic KSHV infection are triggered by the activation of the expression of the Replication and Transcription Activator (RTA). RTA-induced reactivation from latency is followed by immediate-early (IE), and early (E) gene expression, viral genome replication, late (L) gene expression, and finally virion assembly and egress[10]. In a previous study, lytic KSHV endothelial cells treated with 5-tetradecyloxy-2-furoic acid (TOFA), a known allosteric inhibitor of acetyl-CoA carboxylase-α (ACCA) or fatty acid synthesis, showed unaltered viral gene expression and replication. However, both intra- and extracellular infectious virion production was heavily reduced. Transmission electron microscopy (TEM) showed that TOFA-treated cells resulted in oddly shaped virion capsids while nuclear virus factories remained unaffected. It was concluded that inhibition of fatty acid synthesis alters KSHV assembly and egress[11]. Similarly, in murine herpesvirus 68 (MHV-68), a homologous gammaherpesvirus that infects mice, inhibition of lipogenesis by the drug TOFA, and subsequent viral titers during lytic infection, showed that virion production was significantly reduced. Lipidomic analysis of lytic MHV-68-infected cells revealed that lipid remodeling occurs during the early and late phases of infection, resulting in early lipogenesis and triacylglycerol (TG) uptake, and subsequent lipolysis, lipogenesis, or both. There was also a gradual decrease in CEs observed[12]. In another study, in lytic KSHV-infected endothelial cells, 3 to 24 days postinfection, TG content in lipid droplets increased to support virion assembly[6,9]. Another study reported that the attenuation of the cholesterol synthesis pathway in macrophages, via treatment with statins, decreased *in vitro* MHV-68 replication[13]. Another group has reported cholesterol depletion, in iSLK.BAC16 cells, disrupts the integrity of lipid rafts, intracellular signaling and transport micro-domains, which results in significantly reduced virion egress 4.5 days after lytic KSHV induction[14]. These studies reveal that alterations to cholesterol homeostasis occur during the lytic phase of gammaherpesvirus infection. This partly indicates an altered host lipidome and therefore, it is essential to examine and validate any and all alterations to the lipidome during lytic KSHV infection as well. Although some alterations to lipid metabolism during the lytic phase of KSHV infection are reported, the entire lipidome and the molecular mechanisms that drive such alterations remain unknown.

Fatty acid binding proteins (FABPs) are cytosolic proteins that play a central role as lipid chaperones. They traffic long-chain fatty acids to specific cellular compartments for the purpose of modulating lipid metabolism through oxidation, enzyme activity regulation in the cytosol, storage, lipid-mediated transcriptional regulation in the nucleus, paracrine/autocrine signaling outside the cell, and membrane synthesis and signaling in the ER[15–17]. There are twelve known FABPs, ten mammalian (FABP1-9, 12), named and classified based on the tissues they are most abundantly expressed in. Despite their classification, FABPs are not tissue- or cell-specific[18]. A previous study showed that in normoxic and hypoxic KSHV-infected BJAB (a human B cell line) cells, FABP1, FABP4, and FABP7 were significantly upregulated[19]. Similarly, in our KSHV-infected iSLK cell line (iSLK.BAC16), FABP4 and FABP7 were observed to be significantly upregulated during the lytic phase of infection. A study published on Severe Acute Respiratory Syndrome Coronavirus 2 (SARS-CoV-2), an RNA virus, established that FABP4 colocalizes with replication organelles, and inhibition of FABP4 impairs virus replication[20]. In breast cancer, FABP4 and CD36 (fatty acid importer) directly interact for the purpose of regulating lipid metabolism and fatty acid transport into the cell[21].

In this study, we demonstrate that KSHV rewires the host lipidome during lytic KSHV infection and that FABP4 plays a role in KSHV virus production. We report that while early and late lytic gene expression remain unchanged upon FABP4 knockdown, lytic protein translation is reduced. Loss of FABP4 during lytic KSHV infection culminates in a significant reduction in both intracellular and extracellular virus titers, indicating that lytic-induced FABP4 upregulation is required for maximal infectious virion production. Overall, we report the significance of FABP4 during lytic KSHV infection. Through this study, we have uncovered lipid-related metabolic alterations and novel targets of lytic infection, identifying FABP4 as a potential biomarker of KSHV lytic infection.

## Materials and Methods

### Cells and media

The cell lines used in this study are as follows: Inducible SLK cells containing KSHV BAC16 (iSLK.BAC16) and inducible SLK cells (iSLK) used as a control. The iSLK.BAC16 cell line stably maintains the KSHV genome. It is latently infected with recombinant KSHV BAC16, which encodes constitutive expression of GFP. These cells also encode the viral Replication and Transcription Activator (RTA) transgene, which is doxycycline (Dox) and sodium butyrate (NaB) inducible. This triggers the transition in iSLK.BAC16 from latent to lytic phase of infection[22–24]. iSLK cells do not contain the KSHV genome and therefore are used as a control in our experiments[24]. iSLK.BAC16 cells were cultured in DMEM media (Gibco, 1195-065) supplemented with 10% FBS (Corning, 30-249-CR), 10 mM L-Glutamine (Gibco, 25030081), 1µg/mL puromycin (Mirus, 22103916), 250µg/mL G-418 (Fisher Bioreagents, BP673-5) and 1000 µg/mL hygromycin (Corning, 30-240-CR) at 37°C and 5% CO_2_. iSLK cells were cultured in similar media and conditions, but without hygromycin.

### siRNA Knockdown

iSLK and iSLK.BAC16 cells were seeded at 1×10^5^ cells/well onto a 6-well plate. At 24 hours post-seeding, cells were transfected with 25 pmol FABP4 siRNA (siFABP4, GeneGlobe ID: GS2167;) (Qiagen # 1027416) or AllStars negative control siRNA (siASN, Qiagen #1027295) using Lipofectamine® RNAiMAX (Invitrogen #13778-150) in Opti-MEM® Medium (Gibco #11058-021) following manufacturer’s instructions. siRNA-transfection of iSLK and iSLK.BAC16 cells occurred at 37°C and 5% CO_2_ atmosphere for 24 hours. Next, cells were treated with or without Dox and NaB (Dox/NaB) for 48hr. Dox/NaB treatment is required to induce lytic reactivation in iSLK.BAC16 cells.

### Quantitative reverse transcription-PCR

iSLK and iSLK.BAC16 cells were seeded at 1×10^5^ cells/well onto a 6-well plate. The following day, cells were transfected with either siFABP4 or siASN for 24 hours. Next, cells were treated with 1µg/mL of doxycycline (Dox) and 1mM of sodium butyrate (NaB) for 36 hr or 48 hr. After 36/48 hr of Dox/NaB treatment, RNA was extracted using the PureLink^TM^ RNA Mini Kit. Two-step quantitative real-time reverse transcription-PCR (Applied Biosystems) was used to measure the relative mRNA expression of FABP 1-7, ORF59, and ORF26 using the primers listed in **Table 1** in iSLK cells (Dox/NaB -/+) and iSLK.BAC16 cells (Dox/NaB -/+).

### Western Blots

FABP4 knockdown and 48 hour Dox and NaB reactivated iSLK.BAC16 cells were lysed in RIPA buffer, and total protein levels for lysates were quantified using Pierce BCA Protein Assay Kit (ThermoFisherScientific #23225) following the manufacturer’s specifications. For each condition 30µg of total protein lysate was loaded onto 4-20% 10-well Mini-PROTEAN TGX Gel (BIORAD, #4561094) and transferred onto PVDF membranes (BIORAD, #1620264). The membranes were blocked for 1 hour on a rocker with 5% skim milk in a 0.1% TBST solution (1× Tris-buffered saline with 0.1% Tween 20) (Thermo scientific #28360) and probed with primary antibodies diluted in 5% skim milk-TBST by rocking at 4°C overnight. Secondary antibodies were diluted in 5% skim milk-TBST and reacted on a rocker at room temperature for 2hr. Immunolabeled proteins were visualized using Clarity Western ECL substrate and an Amersham ImageQuant™ 800 Western blot imaging system (Cytiva). Immunodetection was performed using mouse antibodies against ORF45 (Sigma Aldrich, #SAB5300153, 1:1000), K8.1 (Millipore, #MABF2678, 1:500), and rabbit antibodies against alpha tubulin (Proteintech # 11224-1-AP. 1:2000). Secondary antibodies included HRP conjugated goat anti-rabbit (Jackson Immuno Research Laboratory, #111-035-003, 1:20,000) or anti-mouse IgG (Jackson Immuno Research Laboratory, #115-035-003, 1:20,000). Western blots were performed in three biological replicates and representative western blot images are included.

### Extracellular virus titer assay

iSLK.BAC16 cells were seeded at 1×10^5^ cells/well onto a 6-well plate. The following day, these cells experienced 24 hr of siASN transfection or FABP4 siRNA-mediated knockdown. Next, after 48 hr of Dox/NaB treatment, cell-free supernatants containing extracellular infectious virus were collected. Supernatants were then centrifuged at 2000 rpm for 10 minutes to remove dead cells and debris. Supernatant titers were introduced onto iSLK cells in a 12-well plate with a confluency of 20,000 cells per well. To each well, 1 μg/ml Polybrene (EMD Millipore #TR-1003-G) was added. Next, these iSLK cells were centrifuged at 300xg for 1 hour and then incubated for 4 hours at 37°C to encourage infection. The cells were then washed with PBS, and fresh media was introduced. To establish latency, cells were further incubated for 48 hours at 37°C and 5% CO_2_.

### Intracellular virus titer assay

Following supernatant collection for the extracellular infectious virus titer experiment, iSLK.BAC16 cells were washed with PBS, trypsinized, and then collected in microcentrifuge tubes. Next, these cells were placed in and out of a −80°C freezer for 3 freeze-thaw cycles for cell lysis. Lysed cells were centrifuged at 2000 rpm for 10 minutes, and then the cell-free supernatant was collected. Titers were then introduced onto iSLK cells in a 12-well plate with a confluency of 20,000 iSLK cells per well, with the addition of 1 μg/ml Polybrene (EMD Millipore #TR-1003-G). Next, the iSLK cells were centrifuged at 300xg for 1 hour and then incubated for 4 hours at 37°C to encourage infection. The cells were then washed with PBS, and fresh media was introduced. To establish latency, cells were further incubated for 48 hours at 37°C and 5% CO_2_.

### Flow Cytometry

Flow cytometry was used to determine the infection rate of KSHV virions based on GFP+ cells. 48 hours after extracellular and intracellular virus titer infection experiments, cells were washed with PBS, collected, and centrifuged at 300xg for 5 minutes. Cell pellets were then resuspended in 100µl FACS buffer (2% FBS in PBS) and then fixed with 4%PFA for 15 minute incubation on ice. Cells were then washed with PBS and centrifuged at 300xg for 5 minutes at 4°C. Cell pellets were then resuspended in 500µl FACS buffer. BD Fortessa SORP flow cytometer, equipped with FITC detection filter, was used to perform flow cytometry. A record of 10,000 cellular events per sample was used during FACS data collection to achieve statistical significance. Data was analyzed by FlowJo V.10.9, in which forward scatter (FSC) and side scatter (SSC) were used to identify relevant cell populations. GFP- and GFP+ were differentiated by quadrant gating, corresponding to FITC- and FITC+ cells, respectively. Statistical significance and bar graph generation were achieved using an unpaired t-test in GraphPad Prism 10.5.0 software. Data is reported as %infection (GFP+) to each sample’s non-induced controls (siASN- and siFABP4-).

### Bulk Lipidomics

In 10cm dishes, iSLK BAC16 cells were seeded and then treated with Dox (1µg/ml) and NaB (1mM) for 48 hours to induce lytic infection. After reactivation, the cells were then washed with PBS, trypsinized, and collected in microcentrifuge tubes. The cell pellets were washed again with sodium bicarbonate and then flash frozen in liquid nitrogen. For each of the two conditions, iSLK.BAC16 cells treated with or without Dox/NaB (Dox/NaB -/+) for 48 hours, 5 biological replicates were generated. The samples were transported to the Northwest Metabolomics Research Center (NW-MRC) for further processing. Lipidomic data were normalized by protein concentration as an internal control. In the web-based platform Metaboanalyst 6.0, lipidomic data were again normalized by log (2) transformation and auto-scaled for further analysis. Statistical analysis and graph making were performed on Metaboanalyst 6.0 and the software GraphPad Prism 10.5.0.

### Statistical Analysis

Figures and statistical analyses were generated using GraphPad Prism 10.5.0 software. Experiments were performed at least in triplicate. Data are presented as the mean ± SEM. Experimental conditions were compared to control conditions using ANOVA or an unpaired t-test. Significant p-values are noted in each figure (* ≤ 0.05; ** ≤ 0.01; *** ≤ 0.001), ns denotes non-significant change.

## RESULTS

### KSHV rewires the host lipidome during lytic infection

Targeted bulk lipidomics was performed on the reporter cell line iSLK.BAC16 to generate a lipid profile of major lipid classes and species as a means of understanding alterations to the host lipid metabolism upon lytic reactivation. The iSLK.BAC16 reporter cell line is a doxycycline-inducible RTA-expressing SLK cell line that stably maintains the bacterial artificial chromosome 16 (BAC16)-KSHV[22]. iSLK.BAC16 cells were seeded and treated with Dox/NaB for 48 hours to induce lytic infection. Cells were then collected and washed with sodium bicarbonate before the pellets were flash frozen. Targeted lipidomic analysis included 5 biological replicates for each condition, normalized to internal protein concentration.

First, we conducted principal component analysis (PCA). The 2D scores plot generated was used to visualize clustering patterns and show clear separation between the samples of each experimental condition according to its principal components[25–27]. The scores plot shows clear separation between our two experimental conditions (**Figure 1A**). Next, we determined the changes to each available lipid class, a broad category based on the lipid head group structure, upon KSHV lytic reactivation[28]. We utilized t-tests to statistically analyze the relative abundances (concentrations of each lipid class normalized by sample protein concentration) of each lipid class in iSLK.BAC16 cells post 48hr in the presence or absence of Dox/NaB. During lytic infection, there is a significant increase in the relative abundance of triacylglycerol (TG), while there is a significant decrease in the relative abundances of cholesterol esters (CE), phosphatidylcholine (PC), and phosphatidylethanolamine (PE) (**Figure 1B, Supplementary Table 1 and 2**). To determine changes to each lipid species, we first normalized raw concentration values by the sample protein concentration, then we used the free web-based software Metaboanalyst to run statistical analysis and generate both a heatmap and a volcano plot. The volcano plot displays the top significant lipid species when comparing Dox/NaB treated over non-treated conditions; the greatest significance, determined by T-tests and fold change analysis, was held by CE species and TGs (**Figure 1C**). The heatmap shows the top 200 significant hits, and the most significant change in Dox/NaB treated samples lies in the relative abundance of CE, TG, PC, and PE (**Figure 1D**). These significant changes to the lipid class and lipid species (based on chain length and bonds) in the host cell indicate that KSHV rewires the host lipidome during lytic infection[29].

**Figure 1.**
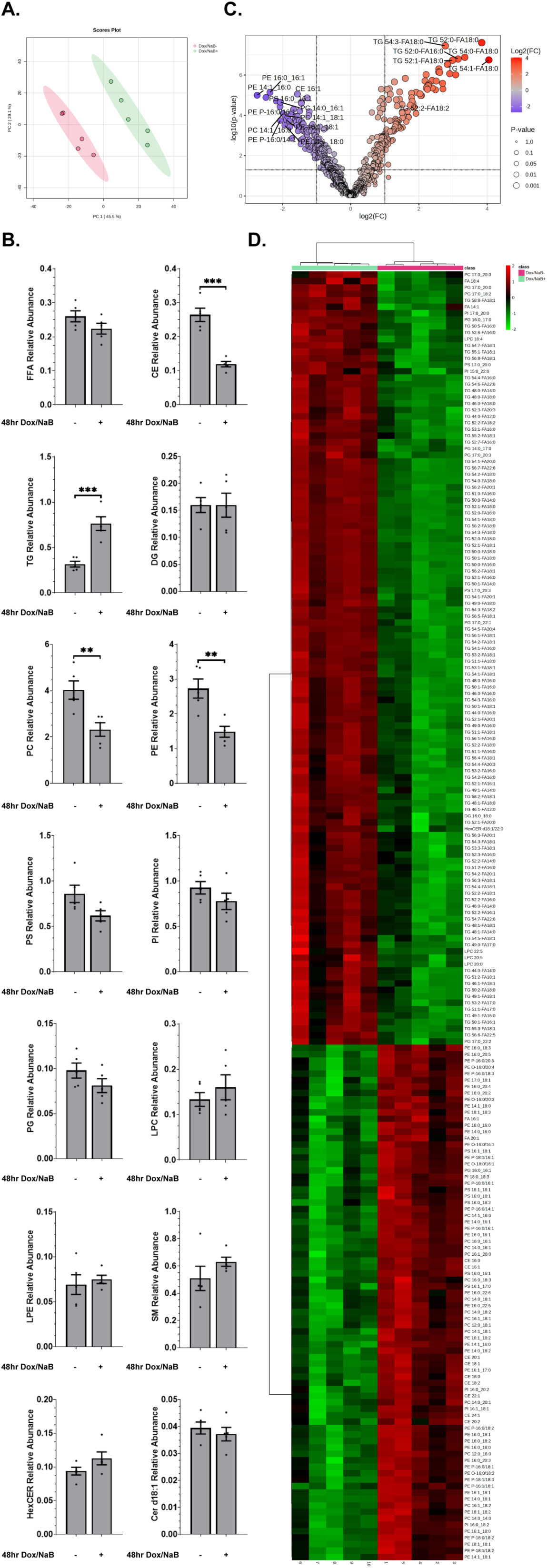
Lipidomic profiling indicates that KSHV rewires the lipidome during lytic infection. Score plots generated via the PCA classification method of Dox/NaB -/+ experimental groups show sample integrity via separation (**A**). Bar graphs show the relative abundance of major lipid classes in latent and 48 hr lytic-induced iSLK.BAC16 cell samples (**B**). The volcano plot shows the top significant lipid species in Dox/NaB+ over Dox/NaB-samples (**C**). A heatmap of the top 200 significant lipid species is displayed and generated via Metaboanalyst software (**D**). ***; P≤0.001. n=5.

### FABP4 is upregulated at the transcript level during lytic KSHV infection

To determine the transcript levels of FABPs during lytic KSHV infection, we measured the relative mRNA expression of FABP1-7 using RT-qPCR. FABP expression was determined in both iSLK.BAC16 cells and iSLK control cells, which lack the KSHV genome, in the presence or absence of Dox/NaB [22–24]. Cells were transfected with either siFABP4 or AllStars Negative siRNA (siASN) as an siRNA control. At 24 hours post-transfection, both iSLK.BAC16 and iSLK cells were treated for 36 or 48 hours with Dox/NaB. Next, RNA was isolated, and RT-qPCR was used to measure the relative gene expression of FABP1-7. FABP4 upregulation is observed at both 36 and 48 hours upon Dox/NaB treatment of iSLK control cells, ∼27.5-fold and 12.5-fold, respectively (**Figure 2A and 2C**). However, the upregulation of FABP4 is much higher in induced iSLK.BAC16 cells at both 36 and 48 hours, ∼75-fold and 38-fold, respectively (**Figure 2B and 2D**). Although FABP5 and FABP7 were seen to be significantly expressed following the lytic reactivation of iSLK.BAC16 cells, there was a significant upregulation of FABP5 and FABP7 in our iSLK control cells treated with Dox/Nab. All other human FABPs either showed no significant change upon Dox/NaB treatment or were very lowly expressed in the experimental cell lines (**Supplementary Figure 1A-1F**). Of the FABP1-7 family, FABP4 transcript levels were the most elevated during lytic KSHV infection.

**Figure 2.**
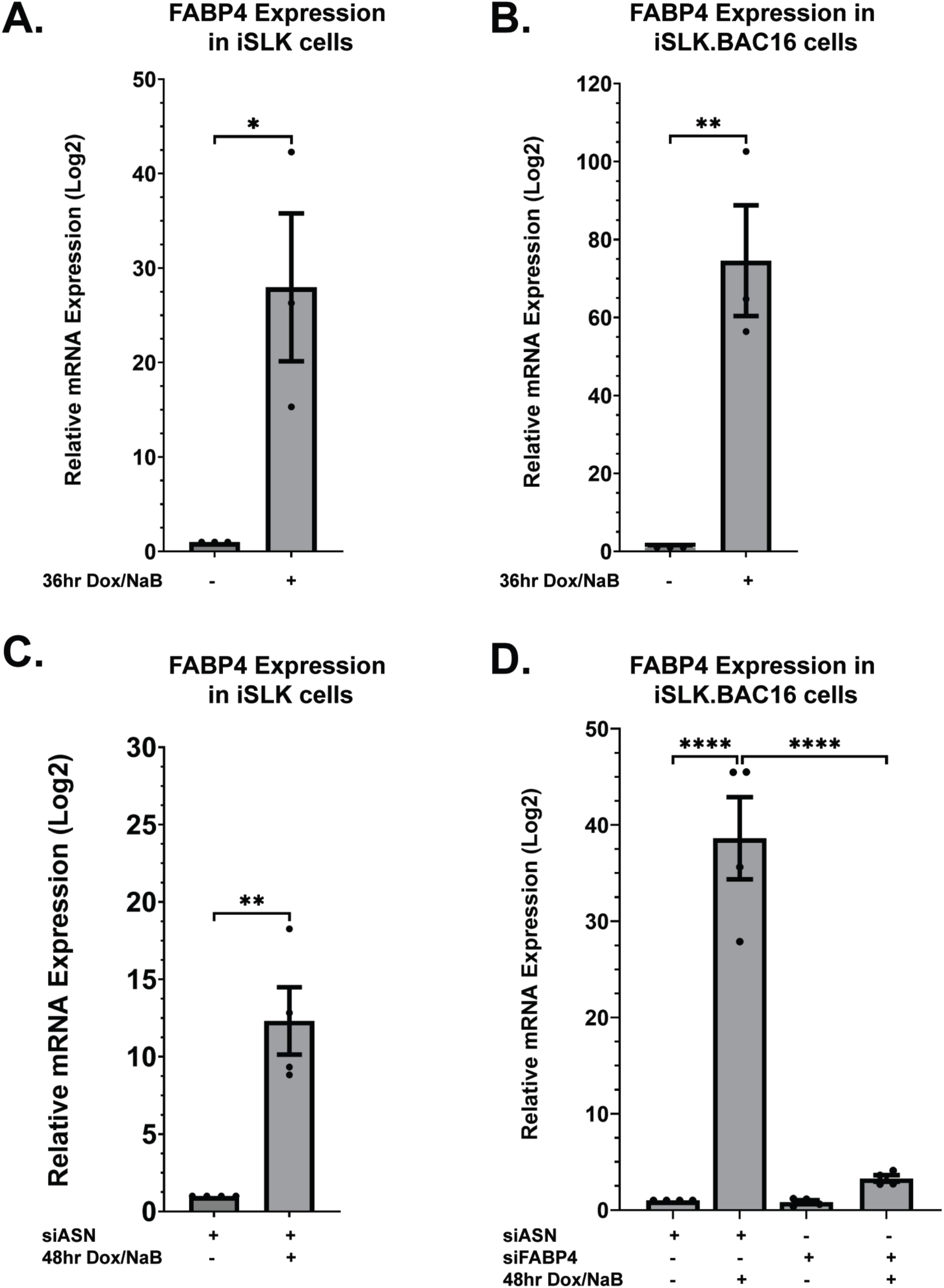
Lytic KSHV infection impacts fatty acid binding protein 4 (FABP4) at the transcript level. Fatty acid binding protein 4 (FABP4) relative mRNA expression levels indicate significant upregulation following 36 hr (**A-B.**) and 48 hr (**C-D.**) of Dox/NaB lytic reactivation in iSLK.BAC16 cells. Successful fatty acid binding protein 4 (FABP4) knockdown to basal relative mRNA expression levels was achieved in 48hr lytic induced iSLK.BAC16 cells (**D.**) siASN= AllStars Negative Control siRNA. ***; P≤0.001. n=4.

To determine the role of FABP4 upregulation during lytic KSHV infection, we first used siRNA to knockdown FABP4 to basal levels. Upon siFABP4 transfection, FABP4 upregulation was significantly reduced by approximately 92% in reactivated iSLK.BAC16 cells (**Figure 2D**). Next, we determined whether FABP4 knockdown during KSHV lytic infection resulted in the upregulation of other FABPs as compensatory expression. We measured the expression of FABP1-7 after siFABP4 treatment in reactivated iSLK.BAC16 cells. No significant upregulation in the expression of FABP1-7 was observed upon FABP4 knockdown (**Supplementary Figure 1A-1F**). Overall, this shows that no compensatory FABP1-7 expression occurs following FABP4 knockdown.

### FABP4 is not required for KSHV gene transcription but is required for viral protein translation

We determined whether FABP4 upregulation is required for early and late viral gene transcription via RT-qPCR (**Figure 3A**). First, iSLK.BAC16 cells were transfected with siASN or siFABP4. At 24 hours post-transfection, cells were induced with Dox/NaB for 48 hours. RNA was isolated, and RT-qPCR was performed to measure specific KSHV genes; the early lytic gene, DNA processivity factor (ORF59), and the late gene, ORF26, which encodes a capsid protein[30,31]. Relative transcriptional expression of ORF59 and ORF26 in iSLK.BAC16 cells transfected with siASN or siFABP4 after 48 hr of mock or Dox/NaB treatment were not affected by FABP4 knockdown during lytic KSHV infection (**Figure 3B-C**). To determine whether FABP4 upregulation is required for early and late viral protein translation we performed western blots in three biological replicates. Upon reactivation after siASN or siFABP4 treatment, iSLK.BAC16 cells were lysed in RIPA buffer to extract protein. Total cell lysates were quantified via BCA assay. Western blot analysis revealed a decrease in the protein expression of ORF45, an early lytic gene encoding a KSHV tegument protein, upon FABP4 knockdown in lytic iSLK.BAC16 cells (**Figure 3D**)[32]. Additionally, the protein level of K8.1, a late lytic gene encoding a KSHV envelope glycoprotein, was decreased in lytic siFABP4 samples (**Figure 3E**)[33]. Here, we have shown that FABP4 does not play a role in the early and late lytic gene expression phase of the KSHV lytic replication lifecycle however, there is a decrease in the protein levels of ORF45 and K8.1.

**Figure 3.**
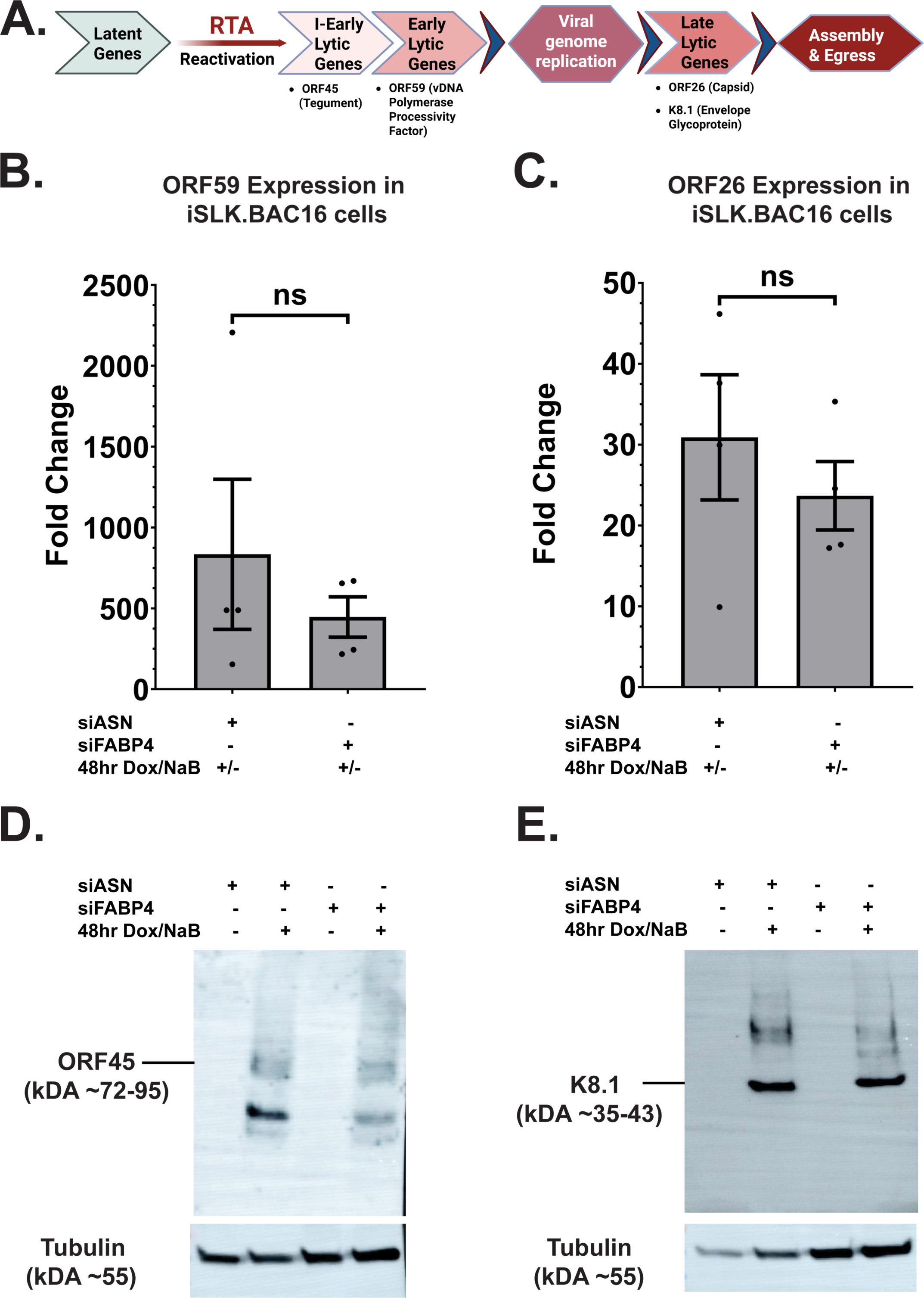
FABP4 upregulation is required for lytic gene translation. An illustrative overview of the lytic cascade, beginning with lytic induction via RTA and ending with virion assembly and egress **(A)**. In FABP4 knockdown and 48hr lytic induced iSLK.BAC16 cells, the relative mRNA expression levels of early (**B.**) and late (**C.**) lytic genes report no significant change. Western Blot analysis shows a decrease in ORF45 protein levels in iSLK.BAC16 cells following siFABP4 and 48hr lytic induction (**D**). K8.1 protein levels are reduced in iSLK.BAC16 cells following siFABP4 and 48hr lytic induction (**E**) siASN = AllStars Negative Control siRNA. ***; P≤0.001. n=4.

### FABP4 upregulation is required for maximal infectious virion production

To determine whether lytic-induced FABP4 upregulation is required for maximal infectious virion production, we performed extracellular virus titer infection assays in triplicate. At 24 hours post siASN or FABP4 knockdown, iSLK.BAC16 cells were treated without, or with Dox/NaB to induce lytic replication. At 48 hours post-induction, cell-free supernatants were collected and titered onto naive iSLK control cells in the presence of 1 µg/ml of polybrene. iSLK cells were then spinoculated for one hour and then incubated in a 37℃ incubator for 4 hours. After incubation, the supernatant was removed, fresh media was added, and cells were further incubated for 48 hours. Cells were then collected and prepared for flow cytometry analysis. We performed flow cytometry to measure the titer of infection, or the number of virus-infected cells based on the mean fluorescent intensity for GFP+ cells using the FITC channel. The representative flow-cytometry histograms show that upon FABP4 knockdown, significantly fewer infected cells are detected in the titer sample (GFP-expressing cells) when compared to the siASN transfection control titer sample. (**Figure 4A**). Upon lytic reactivation, infection with supernatant from siASN-transfected samples resulted in ∼74% GFP+ cells. Meanwhile, infection with supernatant from reactivated and siFABP4 transfected samples resulted in only ∼35% GFP+ cells(**Figure 4B**). These data indicate that knocking down lytic-induced FABP4 upregulation significantly reduces extracellular infectious KSHV virus in iSLK.BAC16 cells.

**Figure 4.**
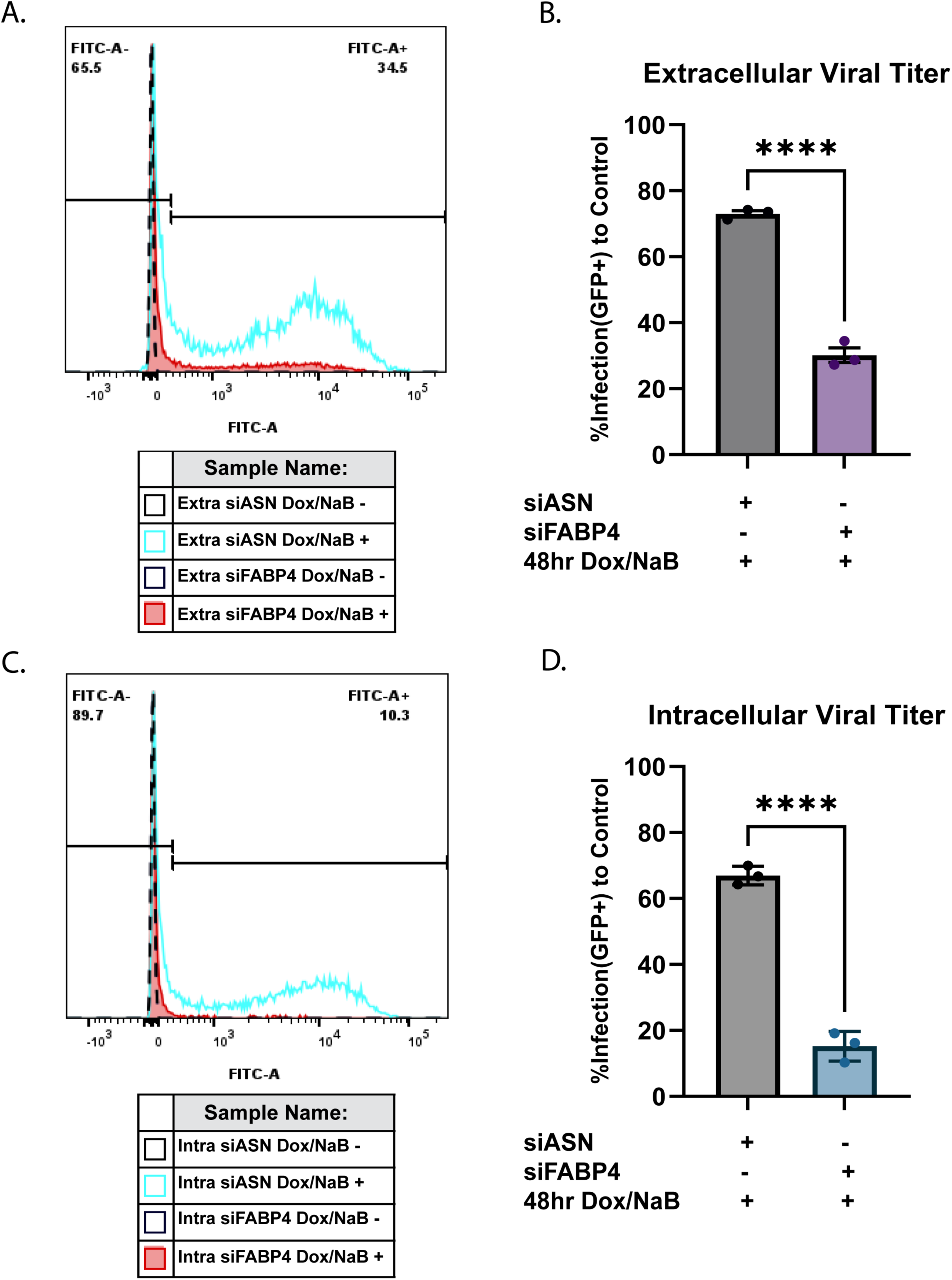
FABP4 is required for successful virion assembly during lytic KSHV infection. Representative flow cytometry-generated images and data graphs of extracellular and intracellular titer infections (**A.**) Representative flow cytometry histogram images of extracellular titer infections compared to control. (**B.**) Relative fluorescence intensity of infected cells to control, quantified via Flow Cytometry, shows a reduction in extracellular titer infection following siFABP4. (**C.**) Representative flow cytometry histogram images of intracellular titer infections compared to control. (**D.**) Relative fluorescence intensity of infected cells to control, quantified via Flow Cytometry, shows a reduction in intracellular titer infection following siFABP4. siASN = AllStars Negative Control siRNA. ***; P≤0.001. n≥3.

To determine whether lytic-induced FABP4 upregulation is required for intracellular infectious virion production, we also performed intracellular virus titer infection assays. After 48 hours of induced lytic-reactivation, siASN or siFABP4-transfected iSLK.BACK16 cells were washed with PBS, exposed to three freeze-thaw cycles for the purpose of cell lysis and intracellular virus extraction. Cell debris was removed via centrifugation, and intracellular virus-containing supernatants were titered onto naive iSLK cells in the presence of 1 µg/ml of polybrene. Cells were then spinoculated for one hour and incubated in a 37℃ incubator for 4 hours. After incubation, the supernatant was removed, fresh media was added, and cells were further incubated for 48 hours before being trypsinized and prepared for flow cytometry analysis. The representative flow-cytometry histograms show that significantly fewer infected cells are detected in the lytic-reactivated siFABP4-transfected titer sample compared to the lytic-reactivated siASN-transfected titer sample (**Figure 4C**).

Upon lytic reactivation, infection with the supernatant from siASN-transfected samples resulted in ∼64% GFP-positive cells. Meanwhile, infection with supernatant from reactivated and siFABP4 transfected samples resulted in only ∼15% GFP+ cells (**Figure 4D**). These data indicates that FABP4 knockdown significantly reduces intracellular infectious KSHV virus upon reactivation of iSLK.BAC16 cells. These data suggest that FABP4 upregulation is contributing to viral assembly and maximal infectious virion production during lytic KSHV infection (**Figure 5**). Overall, loss of FABP4 leads to the reduction of viral protein availability which is a critical downstream requirement for virion assembly and maximal infectious virion production.

**Figure 5.**
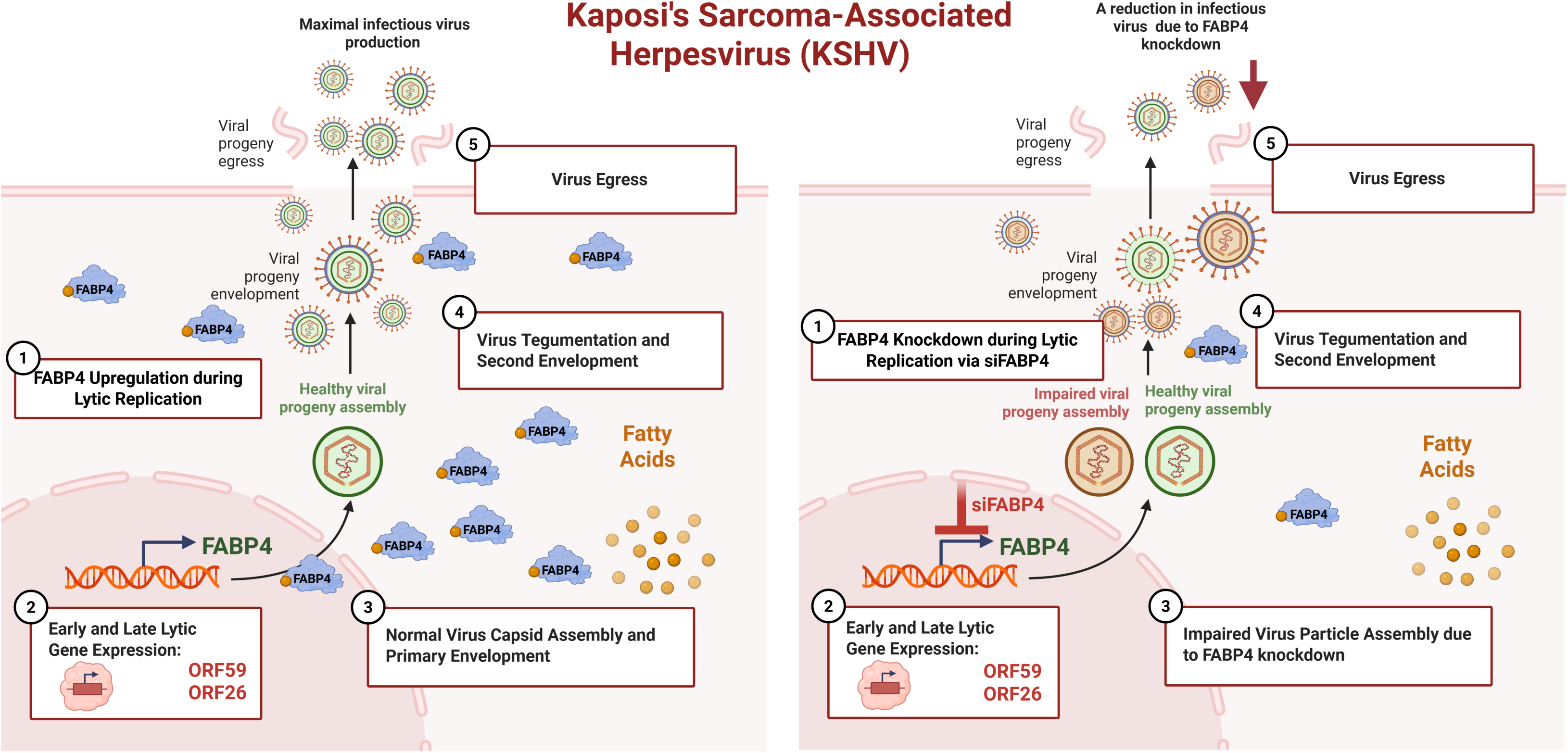
Summary model of the role of FABP4 during lytic KSHV infection. An overview examining how following siRNA-mediated FABP4 knockdown, there is a significant decrease in maximal infectious virion production. Following siFABP4, early and late lytic gene expression remains unaffected. However, impaired viral progeny assembly results in reduced infectious virions and impaired KSHV pathogenesis.

## Discussion

This study determines alterations to the host cell lipidome during lytic KSHV infection and highlights the underlying mechanisms which support it. Previous metabolomic studies in KSHV latently infected endothelial cells have shown a significant increase in lipid abundance compared to mock-infected samples. Furthermore, follow-up functional studies showed that lipogenesis is required for the survival of latently infected endothelial cells[7]. In our study, we used the inducible iSLK.BAC16 system and report a significant increase in the relative abundance of TGs, and a significant decrease in the relative abundances of CE, PC, and PE in lytic KSHV samples compared to latent samples at 48 hours post reactivation (**Figure 1B-1C**). It has been previously reported that TG-enrichment of lipid droplets increases during the lytic phase of KSHV infection in endothelial cells. However, this previous study was reported at very different timepoints than our study, exploring lipid alterations from three to fourteen days post-lytic infection[9]. Interestingly, hepatitis C virus was reported to be dependent on its capsid protein association with TG-rich lipid droplets for virion assembly and infectious virion production[34]. Lipidomic analysis of lytic MHV-68-infected cells revealed that an early (at 8hpi) increase in diacylglycerol (DAG) and TG followed by a TG decrease, free fatty acids (FFA) increase, and a steady state of DAG at 12hpi. These data implicate early lipogenesis and TG uptake, followed by lipogenesis, lipolysis, or both[12]. Our reported significant increase in the relative abundance of TG following lytic induction could be due to TG-rich lipid droplets. During latency, there is an increase in CE synthesis required for microtubule formation and their possible involvement in neo-angiogenesis[9]. Regarding CE metabolism and cholesterol homeostasis, cholesterol is esterified by enzymes Acyl-CoA: cholesterol O-acyltransferases 1 and 2 (ACAT1 and ACAT2) to produce CEs, while CEs are degraded by CE hydrolases to generate fatty acids for β-oxidation and cholesterol for membranes, lipid rafts, and viral envelopes[13,35]. Cholesterol-rich lipid rafts, intracellular signaling and transport micro-domains, are required for successful KSHV virion production. Cholesterol depletion and the inhibition of cholesterol biosynthesis disrupts the integrity of lipid rafts, which results in reduced KSHV virion egress[14]. Therefore, during lytic infection, the decrease in the relative abundance of CE could signify a need for cholesterol in lipid rafts and viral envelopes. Previous lipidomic studies defined the lipid composition of some enveloped viruses, three strains of influenza A virus, and SARS-CoV-2 derived from two different cell types, to include high concentrations of PE and/or PC[36]^-[37]^. The reduction of the relative abundances of PC and PE in our lipidomic data suggest that these lipid classes could be utilized to support the KSHV viral envelope.

To understand the underlying mechanisms which support infectious KSHV virion production, we focused on fatty acid binding proteins (FABPs). Previously, it was reported that in KSHV-infected B-cells (BJAB), FABP1, FABP4, and FABP7 are significantly upregulated under normoxic and hypoxic conditions[19]. Hypoxia is known to induce KSHV productive replication[38]. Following FABP7 knockdown, the KSHV genome copy number was significantly reduced. However, a successful FABP4 knockdown was not generated to investigate the role of FABP4 during KSHV infection[19]. Out of the family of FABP1-7, we report that FABP4 is significantly upregulated in iSLK.BAC16 cells following 36-hour and 48-hour lytic reactivation (**Figure 2, and Supplementary Figure 1)**. While Dox/NaB treatment of iSLK control cells induced FABP4 expression, FABP4 transcript levels were most elevated following Dox/NaB treatment in iSLK.BAC16 cells, suggesting a KSHV-dependent upregulation of FABP4 (**Figure 2**).

FABP4 transcriptional regulation has been linked to the transcription factor, Peroxisome Proliferator-Activated Receptor Gamma (PPARG). PPARG is reported to activate FABP4 expression and FABP4 has been reported to activate the PPARG receptor [39–44]. While the specific mechanism of how FABP4 is upregulated during KSHV lytic infection, we investigated whether any other FABP transcripts could compensate for FABP4 knockdown in this system. Interestingly, FABP4 knockdown in adipocytes induced FABP5 upregulation as a compensatory mechanism[45]. Additionally, the transcriptional profile of latently MHV68-infected mouse spleens identified FABP5 as one of the most differentially expressed genes[46]. However, we report that no significant upregulation of FABP1-7 occurs following FABP4 knockdown during lytic KSHV infection (**Supplementary Figure 1**). These data indicate FABP4 upregulation is driven by KSHV-lytic infection and cannot be compensated by the upregulation of another member of the FABP family.

Moreover, we investigated which stage of lytic KSHV infection requires FABP4 upregulation. The replication and transcription activator (RTA)-induced reactivation begins the lytic cascade of KSHV infection. This is followed by immediate-early (IE), early (E) gene expression, viral genome replication, late (L) gene expression, and virion assembly and egress(**Figure 3A**)[10,47]. We observed that the transcript levels of ORF59, an early lytic gene, showed no significant downregulation in our FABP4 knockdown and lytic-reactivated cells (**Figure 3B**)[31]. Similarly, the transcript levels of ORF26, a late lytic gene, showed no significant downregulation. (**Figure 3C**)[30]. However, we saw a change KSHV lytic protein translation following FABP4 knockdown during lytic replication. The protein levels of ORF45, an immediate early-lytic protein, and K8.1, a late-lytic protein, were significantly decreased following FABP4 knockdown (**Figure 3D-3E**). Therefore, while FABP4 upregulation is not required for lytic gene expression, it is important for lytic protein translation. This requirement potentially stems from the established role of FABP4 in regulating fatty acid oxidation (FAO) and lipolysis, as seen in breast cancer models[21,48–50]. Ultimately, this suggests that KSHV could induce FABP4 upregulation to exploit fatty acid energy reserves, providing metabolic fuel necessary to drive lytic protein translation.

A previous study reported that inhibition of fatty acid synthesis, via treatment with the drug TOFA, in lytic iSLK.BAC16 cells showed unaltered viral gene expression and genome replication. However, both intra- and extracellular infectious virus production was significantly reduced suggesting impaired viral assembly[11]. FABP4 is a cytosolic protein that plays a central role as a lipid chaperone, trafficking fatty acids to specific cellular compartments for various purposes[15–17]. Therefore, we hypothesized that a loss of FABP4 during lytic KSHV infection would result in reduced intracellular and extracellular infectious virus. We report that the loss of FABP4 upregulation results in the reduction of both intracellular and extracellular infectious virus production (**Figure 4**). Since we also observed that FABP4 upregulation is essential for viral protein translation, the reduction in viral protein availability could act as a bottleneck preventing subsequent steps such as viral assembly and maximal infectious virion production.

In conclusion, we have determined that KSHV rewires the host lipidome during lytic KSHV infection. Furthermore, lytic-induced FABP4 upregulation is required for maximal infectious virion production. The specific mechanistic details of FABP4-dependent viral protein translation require further investigation. With this study, we identify lipid-related metabolic vulnerabilities and highlight FABP4 as a novel target of lytic KSHV infection. Our future studies will aim to determine the pharmacological potential of FABP4 inhibition as a strategy to attenuate KSHV pathogenesis.

## Acknowledgements

E.B. and E.L.S. were supported by the National Institutes of Health (NIGMS) 1R35GM160182. We would like to thank all the members of the Sanchez lab, especially Spandan Mukherjee, for their support throughout this study. We acknowledge and thank the Biology Department at the University of Texas at Dallas. We would further like to thank the staff of Northwest Metabolomics Research Center (NW-MRC) and Dr. Zhu Wentao for their aid in lipidomics.

## Key Terms and Abbreviations

KSHV: Kaposi’s Sarcoma Herpesvirus
KS: Kaposi’s Sarcoma
FABPs: Fatty Acid Binding Proteins
TG: Triacylglycerols
CE: Cholesterol Esters
PC: Phosphatidylcholine
PE: Phosphatidylethanolamine

---

**Supplementary Figure 1.** FABP regulation during lytic KSHV infection FABP1, 2, 3, 5, 6, and 7 relative mRNA expression levels following 48hr of Dox and NaB treatment reveal that only FABP7 and FABP5 are significantly upregulated during lytic KSHV infection. FABP1, 2, 3, and 6 showed no significant change in their relative mRNA expression levels in FABP4 knockdown and 48 hr lytic-induced iSLK.BAC16 cells (**E-F**). siASN = AllStars Negative Control siRNA. ***; P≤0.001. n=4.

