## Supplementary figures and images for "Global Lipidomic Analysis of Lytic KSHV Infection: The lipid chaperone FABP4 is required for maximal infectious virion production"

### Supplementary Figure 1

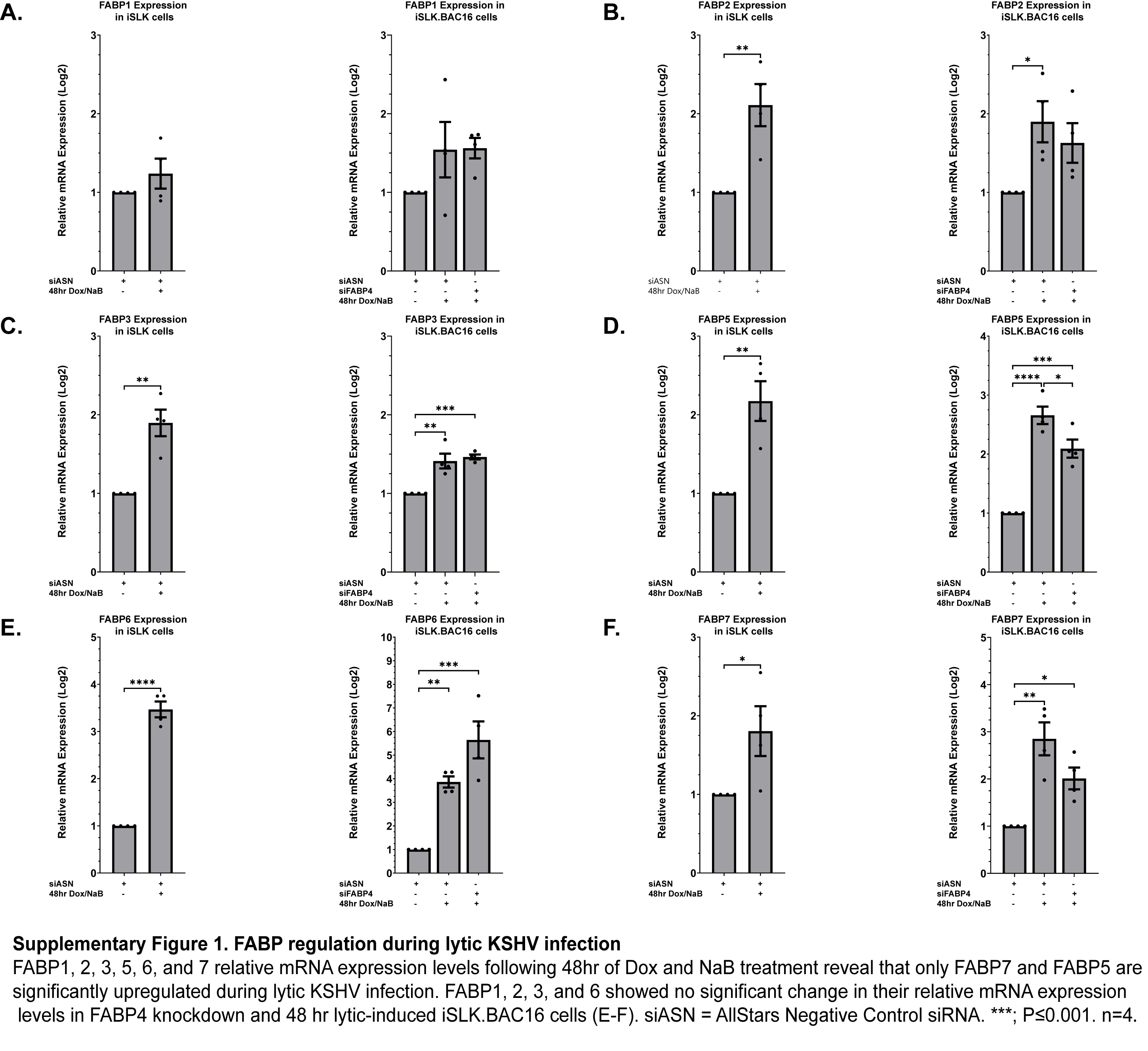
